# Rumen transfaunation between low- and high-methane-yielding dairy cows reveals asymmetric microbiome reconstitution patterns: a pilot study

**DOI:** 10.64898/2026.04.07.716947

**Authors:** Puchun Niu, Carl Kobel, Velma T. E. Aho, Clementina Alvarez, Egil Prestløkken, Peter Lund, Adrian Omar Maynez-Perez, Phil B. Pope, Angela Schwarm

**Affiliations:** Department of Animal and Aquacultural Sciences, Norwegian University of Life Sciences, PO Box 5003, 1432 Ås, Norway; Department of Molecular Biology and Genetics, Plant Molecular Biology, Aarhus University, Aarhus, Denmark; Yara International ASA, 0213 Oslo, Norway; Department of Animal Science, AU Foulum, Aarhus University, 8830 Tjele, Denmark; Centre for Microbiome Research, School of Biomedical Sciences, Queensland University of Technology (QUT), Translational Research Institute, Woolloongabba, Queensland, Australia; Faculty of Chemistry, Biotechnology and Food Science, Norwegian University of Life Sciences, 1432 Ås, Norway

**Keywords:** Methane yield, rumen transfaunation, GreenFeed system, metagenomic and metaproteomic analysis, ruminant

## Abstract

**Background:** This study investigated rumen microbiome reconstitution and methane (CH_4_) emissions following a complete exchange of rumen contents between low- and high-CH_4_-yielding Norwegian Red dairy cows. Twenty cows were screened for CH_4_ yield, and two low and two high emitters were selected for rumen cannulation and content swap. Total rumen contents were swapped after complete evacuation and washing of both the rumen and omasum. Rumen samples were collected twice in weeks −1, 1, 3, and 7 for fermentation analysis, metagenomics, and metaproteomics, and at week 8 CH_4_ production was measured.

**Results:** Prior to the swap, low and high emitters produced 21.2 ± 0.7 and 26.3 ± 1.4 g CH_4_/kg dry-matter intake (DMI), respectively. Eight weeks after swap, CH_4_ yields were 12.7 ± 0.3 and 28.9 ± 0.3 g CH_4_/kg DMI, respectively, showing that the CH_4_ phenotype of each cow was maintained. Analysis of metagenome-derived 16S rRNA gene sequences showed that low emitters gradually re-established their original microbial community, whereas high emitters retained donor-like microbiota. Metaproteomic mapping suggested higher expression of Prevotella-associated succinate–propionate pathway enzymes in low emitters at week 7, though these differences were modest.

**Conclusion:** These findings suggest that host factors influence CH_4_ output and microbial reconstitution, with low emitters restoring their native microbiome while high emitters retained a donor-associated community yet continued to emit high CH_4_. Results should be interpreted with caution given the small sample size (n = 2 per phenotype) and require confirmation in larger studies.

**Importance:** Reducing enteric methane from cattle requires understanding whether the rumen microbiome or the host animal is the primary driver of methane output. We exchanged the entire rumen contents between low- and high-methane-yielding dairy cows and measured methane production alongside metagenomic and metaproteomic profiling over two months. Despite receiving each other’s microbiomes, each cow’s methane phenotype persisted—low emitters stayed low and high emitters stayed high. Microbiome reconstitution was asymmetric: low emitters restored their original microbial community, while high emitters retained the donor microbiota. Methanogen communities did not differ between phenotypes, pointing to host-level rather than microbial-level control of methane yield. These pilot findings suggest that breeding for favorable host traits may be essential for lasting methane reduction, and that microbiome transfer alone is unlikely to shift an animal’s methane phenotype. Larger studies are needed to confirm these observations.

## Introduction

A promising measure to reduce enteric methane (CH_4_) production from ruminant livestock is the manipulation of the rumen microbiome [1]. Basic knowledge of the underlying factors that influence microbial community composition is required to achieve a predictable and consistent reduction in enteric CH_4_ production when manipulating these communities. In addition to the rumen microbiome, host genetics can influence CH_4_ production, but the two have been shown to be largely independent [2]. Hence, breeding and microbial manipulation can offer separate but complementary approaches to CH_4_ mitigation.

Rumen transfaunation, defined as the transfer of rumen content from a healthy donor to a recipient animal, is a long-established veterinary practice to restore rumen function after digestive disorders [3]. Clinical transfaunation restores rumen function in sick animals, whereas experimental rumen content swap is conducted between healthy animals. An early study demonstrated that the rumen microbial community displays strong host specificity and resilience after approximately 95% of the rumen contents were swapped between cows (n = 4) [4]. Subsequent experiments confirmed that the re-establishment of microbial communities after such interventions is highly individualized [5] and that the methanogenic community tends to be more stable than the bacterial fraction following complete transfers [6]. These studies collectively suggest that while the rumen microbiome can be transiently modified, it often reverts, at least partially, toward a host-specific equilibrium.

Recent advances in multi-omics technologies provide opportunities to link rumen microbial composition with its functional activity in vivo. Shotgun metagenomics, particularly when combining short- and long-read sequencing, enables the reconstruction of metagenome-assembled genomes (MAGs) and offers insight into the metabolic potential of individual rumen taxa [7, 8]. Metaproteomics adds complementary information by identifying the proteins expressed in the rumen environment, thereby revealing active metabolic pathways, including those relevant for hydrogen turnover and carbohydrate degradation [9].

However, none of these earlier investigations combined complete rumen content swap between low- and high-methane-emitting cows with direct measurements of enteric CH_4_ emissions and accompanying metagenomic and metaproteomic analyses. Existing studies have characterized microbial succession and fermentation profiles following rumen transfaunation, but CH_4_ production was either not measured or only inferred from fermentation end products.

Consequently, the effect of a full rumen content swap on both microbial reconstitution dynamics and CH_4_ yield (g CH_4_ per kg dry matter intake (DMI)) in lactating dairy cows remains unknown.

Therefore, the objective of the current study was (i) to identify and cannulate lactating Norwegian Red dairy cows characterized by low and high CH_4_ yield, and (ii) to examine microbial and functional community dynamics at early, middle, and late stages after complete swap of rumen content between the two CH_4_ phenotypes using metagenomic and metaproteomic analyses, along with associated ruminal fermentation parameters. Methane production was measured prior to cannulation and again two months after the swap to evaluate whether the CH_4_ phenotype (low or high) of each cow was altered. We hypothesized that the microbial community in the rumen would gradually reconstitute after rumen transfaunation, and the methane category of the cows would remain unchanged, reflecting the dominant influence of host factors over microbial inheritance.

## Results

To establish baseline phenotypic variation, we first characterized feed intake, milk production, body weight, and CH_4_ emissions in 20 Norwegian Red cows during a 30-day screening period. This variation enabled identification of two consistently low and two consistently high emitters for the subsequent rumen content swap experiment. Table 1 summarizes the performance traits and CH_4_ measures across the screened herd, which showed substantial natural variation in DMI, milk yield, and CH_4_ production. Dry matter intake ranged from 16.7 to 28.2 kg/day, milk yield from 21.8 to 51.8 kg/day, and BW from 522 to 782 kg. Methane emissions ranged from 319 to 596 g/day, corresponding to CH_4_ yields of 13.7 to 27.3 g/kg DMI.

**Table 1.** Descriptive statistics of 20 intact cows screened for performance and methane (CH_4_) production during a 30-d pretrial.

| Item | Min | Max | Mean (SD) | CV (%) |
| --- | --- | --- | --- | --- |
| Dry matter intake (DMI), kg/d | 16.7 | 28.2 | 22.2 (2.7) | 12.2 |
| Milk yield (MY), kg/d | 21.8 | 51.8 | 34.8 (7.7) | 22.1 |
| Body weight (BW), kg | 522 | 782 | 611 (66) | 10.8 |
| CH<sub>4</sub> emission |  |  |  |  |
| CH<sub>4</sub>, g/d | 319 | 596 | 475 (68) | 14.3 |
| CH<sub>4</sub>/DMI, g/kg | 13.7 | 27.3 | 21.6 (3.3) | 15.3 |
| CH<sub>4</sub>/MY, g/kg | 7.5 | 19.7 | 14.2 (3.2) | 22.5 |
| CH<sub>4</sub>/BW<sup>0.75</sup>, g/kg | 2.82 | 5.25 | 3.89 (0.63) | 16.2 |

Rumen contents from the 119 four selected cows differed modestly in total mass and particle–liquid composition (Table 2). Particle mass ranged from 53.9 to 67.9 kg fresh matter and liquid mass from 27.3 to 37.5 kg fresh matter, resulting in particle-to-liquid ratios between 1.56 and 2.49. Total rumen content DM ranged from 10.6 to 12.9 kg (1.84–2.11% of BW), confirming that rumen evacuation and subsequent washing were performed under comparable physiological states across animals.

**Table 2.** Characteristics of the rumen content in low- (L1, L2) and high (H1, H2) methane-yielding cows on the day of rumen content swap.

| Animal | Particles, kg FM | Liquid, kg FM | Particle to liquid ratio | Content, kg DM | Content, % DM | Content DM, % BW | Liquid, % BW |
| --- | --- | --- | --- | --- | --- | --- | --- |
| L1 | 58.5 | 37.5 | 1.56 | 11.9 | 12.3 | 1.84 | 5.79 |
| L2 | 53.9 | 27.5 | 1.96 | 11.3 | 13.9 | 1.98 | 4.82 |
| H1 | 67.9 | 27.3 | 2.49 | 12.9 | 13.5 | 2.11 | 4.47 |
| H2 | 55.2 | 27.9 | 1.98 | 10.6 | 12.7 | 1.84 | 4.84 |

FM, fresh matter; DM, dry matter; BW, body weight.
Note: The rumen content was swapped between L1 and H1, and L2 and H2.

The feed intake, milk yield, and CH₄ production of the four cows selected for the rumen content swap were monitored across the first (pre-swap) and second (post-swap) lactations, with the corresponding values summarized in Table 3. Methane measurements were obtained during the screening period before cannulation and again approximately two months after the rumen content swap. Before the swap, low emitters (L1 and L2) produced 467 and 389 g CH_4_/day, corresponding to CH_4_ yields of 20.7 and 21.6 g CH_4_/kg DMI, respectively. High emitters (H1 and H2) emitted 485 and 531 g CH_4_/day, with corresponding yields of 25.3 and 27.5 g CH_4_/kg DMI. Milk yield and DMI were similar between pre- and post-swap periods. At the end of the experiment, CH_4_ yields of the low emitters were markedly lower (12.9 and 12.5 g CH_4_/kg DMI for L1 and L2, respectively), while those of the high emitters remained high (28.7 and 29.1 g CH_4_/kg DMI for H1 and H2). The same trend was observed for CH_4_ intensity (g CH_4_/kg milk yield). These results indicate that, despite the microbial swap, the original CH_4_ emission phenotype of each cow was maintained.

**Table 3.** Performance and methane production of the low-(L1, L2) and high (H1, H2) methane-yielding cows.

| Item | L1 |  | L2 |  | H1 |  | H2 |  |
| --- | --- | --- | --- | --- | --- | --- | --- | --- |
| Measurement period<sup>1</sup> | Pre-swap | Post-swap | Pre-swap | Post-swap | Pre-swap | Post-swap | Pre-swap | Post-swap |
| DMI, kg/d | 22.7<br>(1.7) | 23.4<br>(2.6) | 17.9<br>(1.7) | 20.2<br>(2.2) | 19.0<br>(1.3) | 19.6<br>(3.0) | 19.2<br>(1.7) | 19.8<br>(1.8) |
| MY, kg/d | 37.1<br>(7.4) | 27.1<br>(4.6) | 28.0<br>(4.7) | 31.6<br>(5.8) | 37.0<br>(8.7) | 31.3<br>(7.5) | 28.6<br>(5.5) | 33.3<br>(9.8) |
| BW, kg | 582 | 671 | 522 | 599 | 560 | 614 | 548 | 601 |
| CH<sub>4</sub>, g/d | 467<br>(44) | 298<br>(40.9) | 389<br>(25) | 249<br>(71.6) | 485<br>(46) | 552<br>(47.8) | 531<br>(22) | 571<br>(64.9) |
| CH<sub>4</sub>/DMI, g/kg | 20.7<br>(2.4) | 12.9<br>(2.2) | 21.6<br>(2.5) | 12.5<br>(4.1) | 25.3<br>(2.3) | 28.7<br>(6.3) | 27.5<br>(3.1) | 29.1<br>(4.6) |
| CH<sub>4</sub>/MY, g/kg | 13.1<br>(2.8) | 8.2<br>(2.4) | 14.7<br>(2.6) | 7.6<br>(2.4) | 17.7<br>(2.5) | 18.0<br>(7.6) | 17.4<br>(3.4) | 17.3<br>(4.7) |
| CH<sub>4</sub>/BW<sup>0.75</sup>, g/kg | 3.94 | 2.26 | 3.56 | 2.05 | 4.21 | 4.48 | 4.69 | 4.7 |

<sup>1</sup>Methane was measured during the screening phase in the 1st lactation (pre-rumen swap, n = 30 days) and again at the end of the rumen swap experiment in the 2nd lactation (post-rumen swap, two months after the swap, n = 7 days).
DMI, dry matter intake; MY, milk yield; BW; body weight; CH<sub>4</sub>, methane.
Data are presented as arithmetic means with standard deviations in brackets, except for BW and CH<sub>4</sub>/BW<sup>0.75</sup>, for which BW was measured once in each period.

We next examined bacterial and archaeal community dynamics before and after the swap. Microbial community composition is shown in Figure 1, based on 16S rRNA gene sequences extracted from the assembled metagenomes. Because these sequences were derived from metagenome assemblies rather than targeted amplicon sequencing, coverage may vary across samples, and the resulting profiles should be viewed as indicative of broad compositional trends. The principal component analysis (PCA) of all samples (Fig. 1A) showed that microbial profiles clustered by the time point of sampling. Prior to the swap (week −1), the low emitters (L1 and L2) clustered closely, while the two high emitters (H1 and H2) were widely separated. Following the rumen content swap, high emitters (H1 and H2) that received rumen contents from low emitters (L1 and L2) exhibited microbial community structures closely resembling their respective donor cows throughout weeks 1, 3, and 7 (Fig. 1B,D). In contrast, the low emitters (L1 and L2), which received rumen contents from high emitters, showed a gradual re-establishment of their pre-swap microbial community composition over the same period (Fig. 1C,E). Methanogenic archaea were also examined from the metagenome-derived 16S rRNA gene data. No significant differences in methanogen relative abundance or community composition were observed between low- and high-methane-yielding cows at any time point (data not shown).

**Fig. 1.**
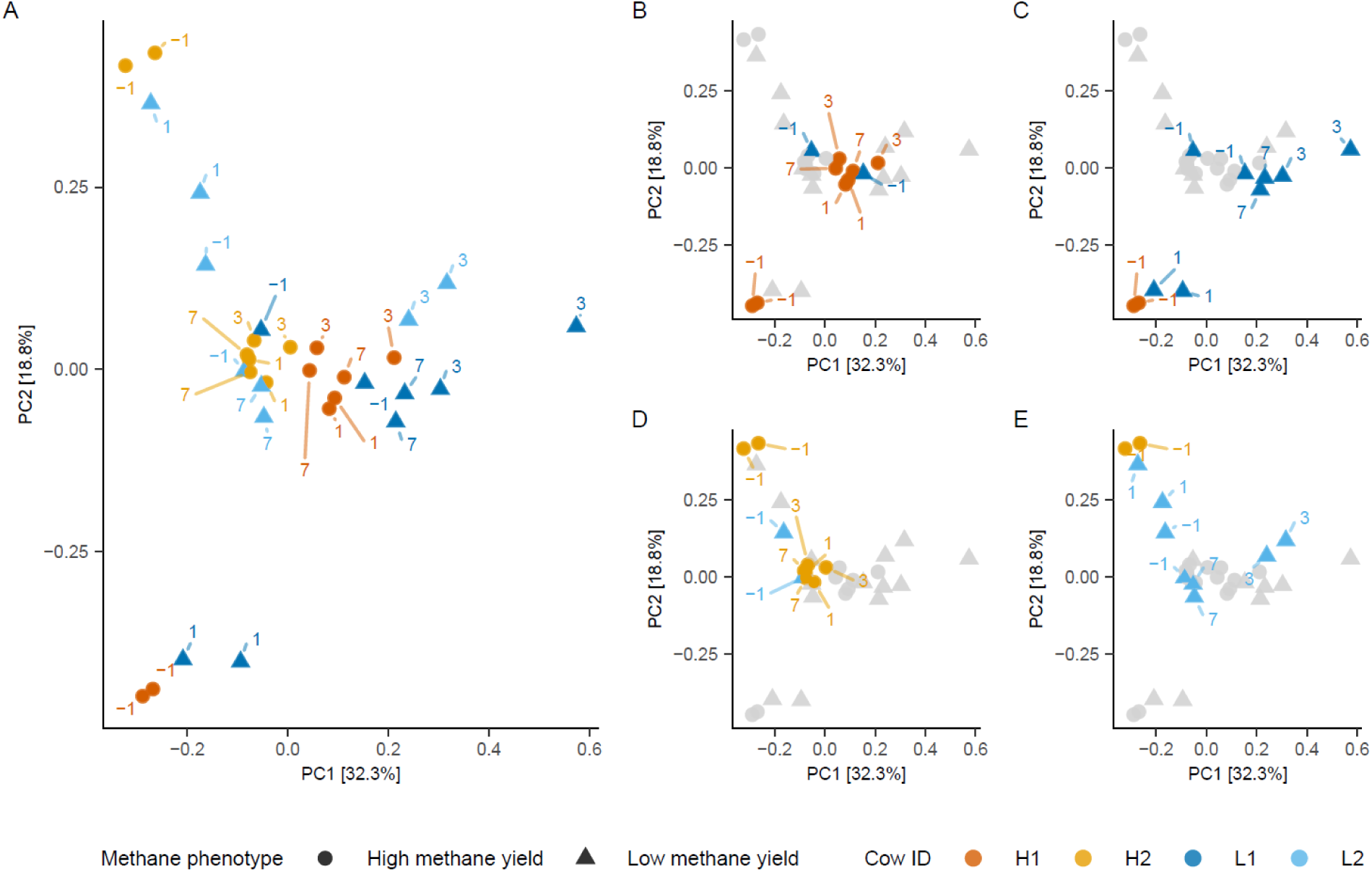
Microbial community composition in low- and high-methane-yielding cows before and after rumen content swap. Methane phenotype: High methane yield and Low methane yield cows (expressed as g CH_4_/kg dry matter intake). Cow IDs: L1 and L2 = low-methane-yielding cow 1 and cow 2; H1 and H2 = high-methane-yielding cow 1 and cow 2. Each dot (●) or triangle (▲) represents the microbial community in a specific sample, with duplicate samples collected each week. Numbers next to points indicate sampling week: –1 (week before rumen swap) and 1, 3, 7 (weeks after rumen swap). (A) Overall microbial composition of all samples from low and high methane-yielding cows across all weeks. (B) Microbial community in cow H1 before the rumen swap (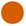, –1) and after receiving rumen content from cow L1 (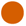, 1, 3, and 7); also shows cow L1 before the swap (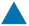, –1). (C) Microbial community in cow L1 before the rumen swap (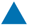, –1) and after receiving rumen content from cow H1 (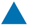, 1, 3, and 7); also shows cow H1 before the swap (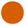, –1). (D) Microbial community in cow H2 before the rumen swap (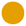, –1) and after receiving rumen content from cow L2 (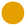, 1, 3, and 7); also shows cow L2 before the swap (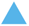, –1). (E) Microbial community in cow L2 before the rumen swap (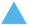, –1) and after receiving rumen content from cow H2 (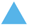, 1, 3, and 7); also shows cow H2 before the swap (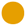, –1).

MAG-based metagenomics and the accompanying metaproteomic profiles provided complementary insight into both the ecological structure and functional activity of the rumen microbiome. In both MAG abundance data, which reflects taxonomic composition (Fig. 2A), and in metaproteomic data, offering a functional view (Fig. 2B), microbial profiles clustered primarily by cow and CH_4_ phenotype rather than by sampling week. Before the rumen content swap (week −1), distinct separation was observed between low- and high-methane emitters. Following the swap, the microbial communities of all cows showed dynamic but individualized temporal shifts through weeks 1, 3, and 7. Despite these changes, no convergence between low-and high-emitter animals was observed, indicating that overall community structure remained phenotype-specific throughout the recovery period.

**Fig. 2.**
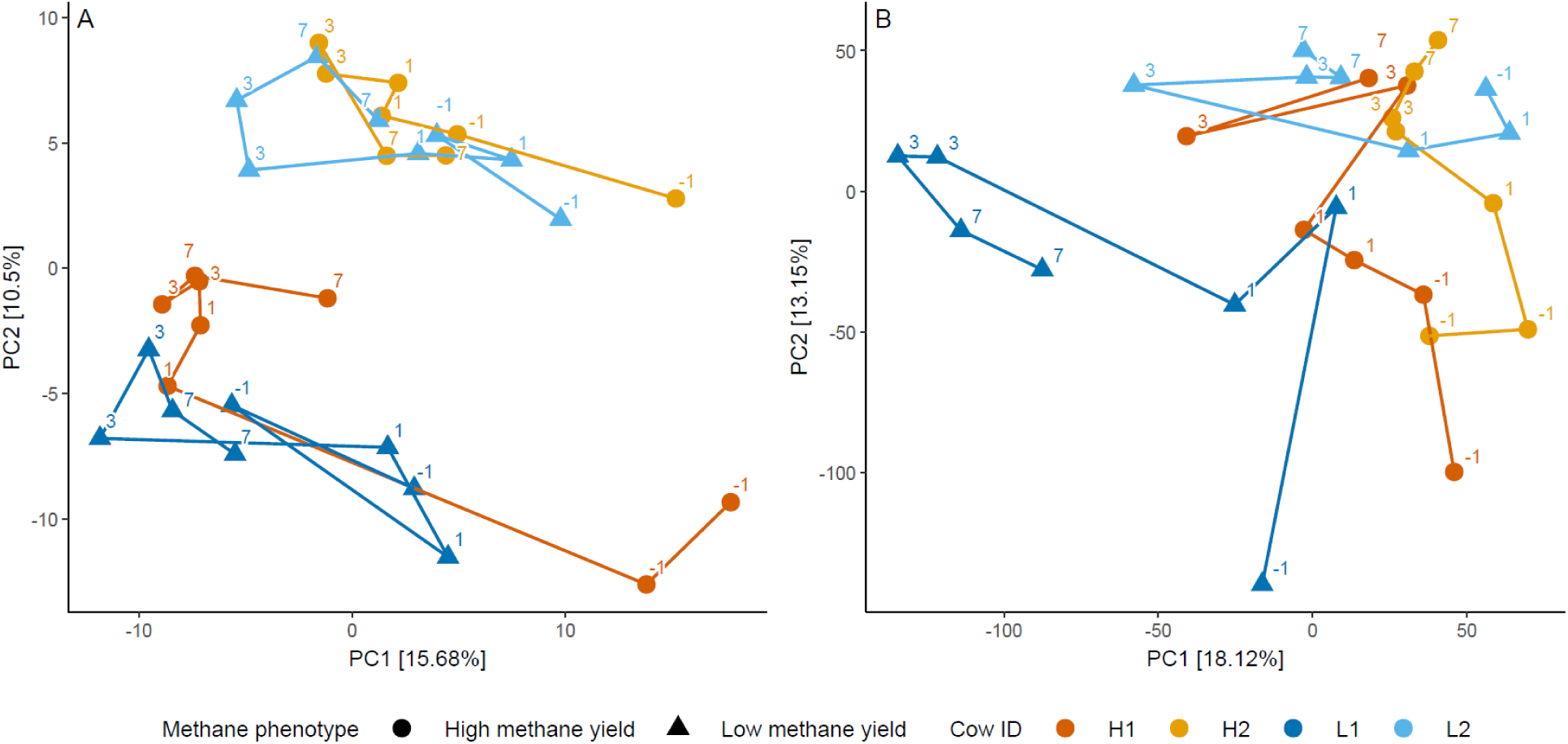
Trajectory of microbial community composition before and after rumen content swap in low- and high-methane-yielding cows (expressed as g CH_4_/kg dry matter intake). Panel A: Microbial community composition based on metagenome-assembled genome (MAG) data. Panel B: Microbial community composition based on metaproteomic data. Methane phenotype: High methane yield and Low methane yield cows (expressed as g CH_4_/kg dry matter intake). Cow IDs: L1 and L2 = low-methane-yielding cow 1 and cow 2; H1 and H2 = high-methane-yielding cow 1 and cow 2. Each dot (●) or triangle (▲) represents the microbial community in a specific sample, with duplicate samples collected each week. Numbers next to points indicate sampling week: –1 (week before rumen swap) and 1, 3, 7 (weeks after rumen swap). Lines connect samples in chronological order. Both panels show trajectories of microbial communities across time for all cows. Axes represent principal component analysis (PCA): MAGs data (PC1 = 15.68%, PC2 = 10.5%) and metaproteomic data (PC1 = 18.12%, PC2 = 13.15%).

Fermentation patterns also exhibited phenotype-specific behavior over time (Fig. 3). Acetate concentrations increased in both CH_4_ phenotypes after the rumen content swap, while propionate decreased slightly over the study period, with numerically higher values in low emitters, though the absolute differences were small (approximately 1.5–2 mol/100 mol). As a result, the acetate-to-propionate ratio was higher in high emitters throughout the experiment. Ruminal pH and ammonia were relatively stable over time within each group, indicating that overall fermentation patterns differed between CH_4_ phenotypes but were not strongly altered by the content swap.

**Fig. 3.**
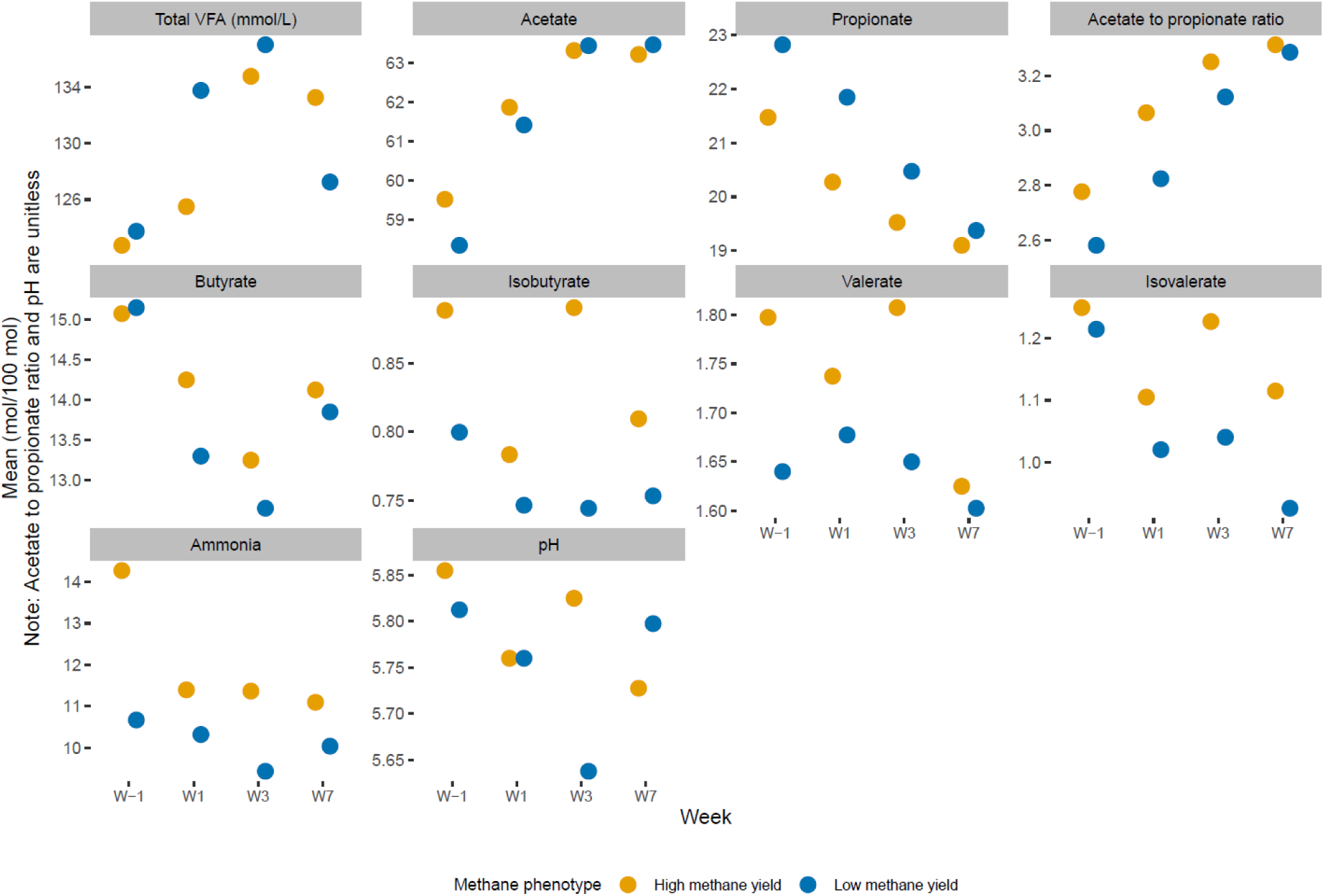
Time course of mean concentrations of total volatile fatty acids (VFAs), individual VFAs, and pH in rumen contents of low- and high-methane-yielding cows (expressed as g CH_4_/kg dry matter intake) at weeks (W) –1, 1, 3, and 7 relative to the rumen swap.

To identify bacterial and archaeal taxa contributing to phenotype-specific functional differences, we quantified MAG-resolved protein intensities (summed LFQ intensity) across weeks (Fig. 4). At week −1, differences in MAG abundance between CH_4_ phenotypes were small, but the contrast widened by week 7. The displayed MAGs represent those showing significant differences (*P* < 0.05) in week 7. Several *Prevotella* species increased in abundance over time in both phenotypes, with overall protein intensities consistently higher in low emitters than in high emitters. A comparable but modest trend was also seen for a few *Isotricha* spp., which showed slightly higher protein intensities in low emitters at week 7 (Fig. S1).

**Fig. 4.**
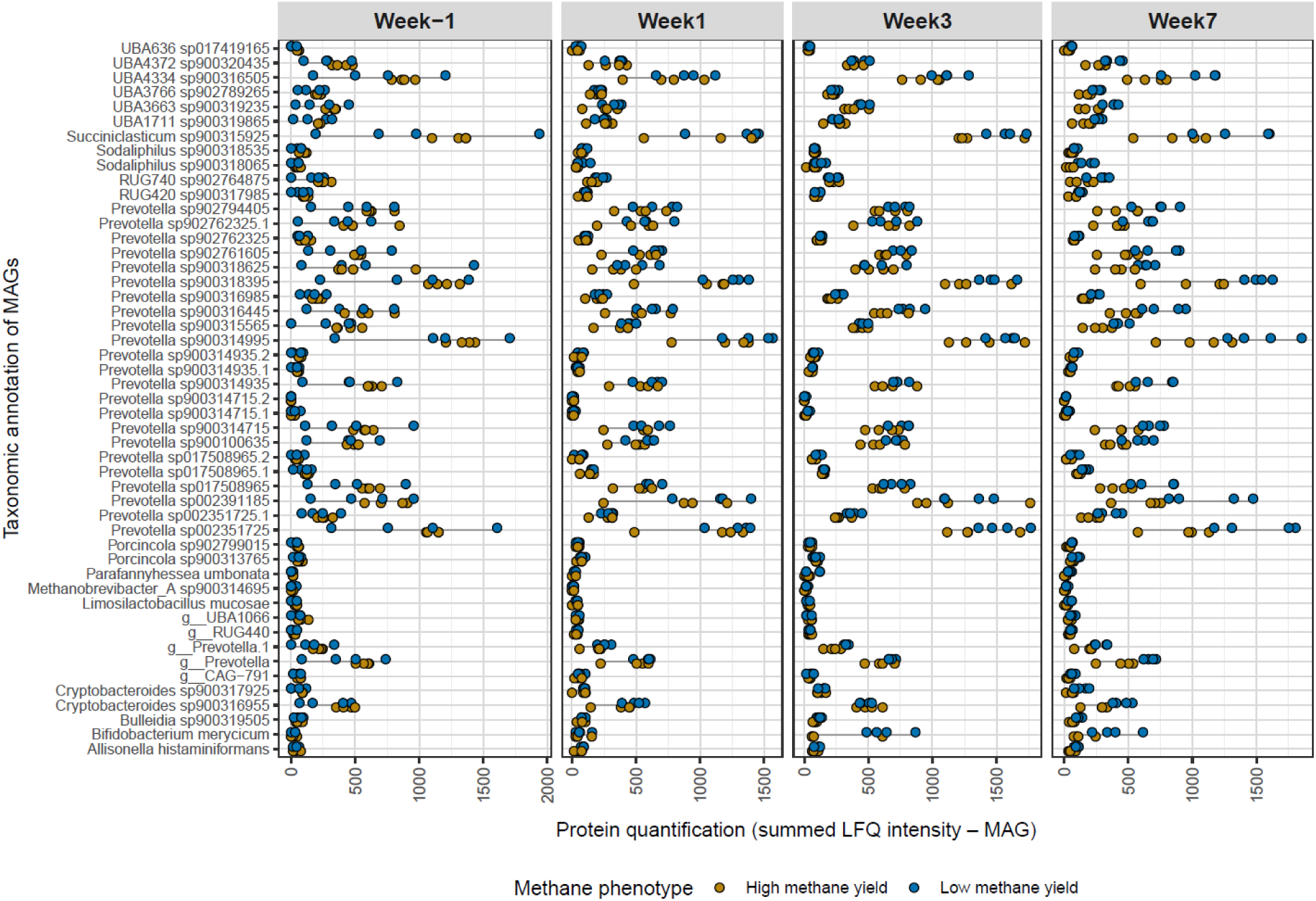
Protein quantification of metagenome-assembled genomes (MAGs; n=49) across experimental weeks for low- and high-methane-yielding cows (expressed as g CH_4_/kg dry matter intake). The MAGs displayed were selected based on t-tests conducted in week 7 using uncorrected p-values (p < 0.05) to highlight taxa showing the strongest phenotype differences. Each point represents a single sample (two cows per phenotype, each sampled twice in each week). Protein quantification values represent the summed LFQ intensities of all detected proteins within each MAG.

Pathway-level protein detection (LFQ, log2) revealed marked differences in metabolic activity across phenotypes and weeks (Fig. 5). Five representative MAGs, including *Prevotella bryantii*, *Prevotella sp.* 900314935, *Prevotella sp.* 900316445, *Prevotella sp.* 900318395, and UBA4334 *sp.* 900316505, are depicted in Figure 5 across weeks −1, 1, 3, and 7 for both CH_4_ phenotypes.

**Fig. 5.**
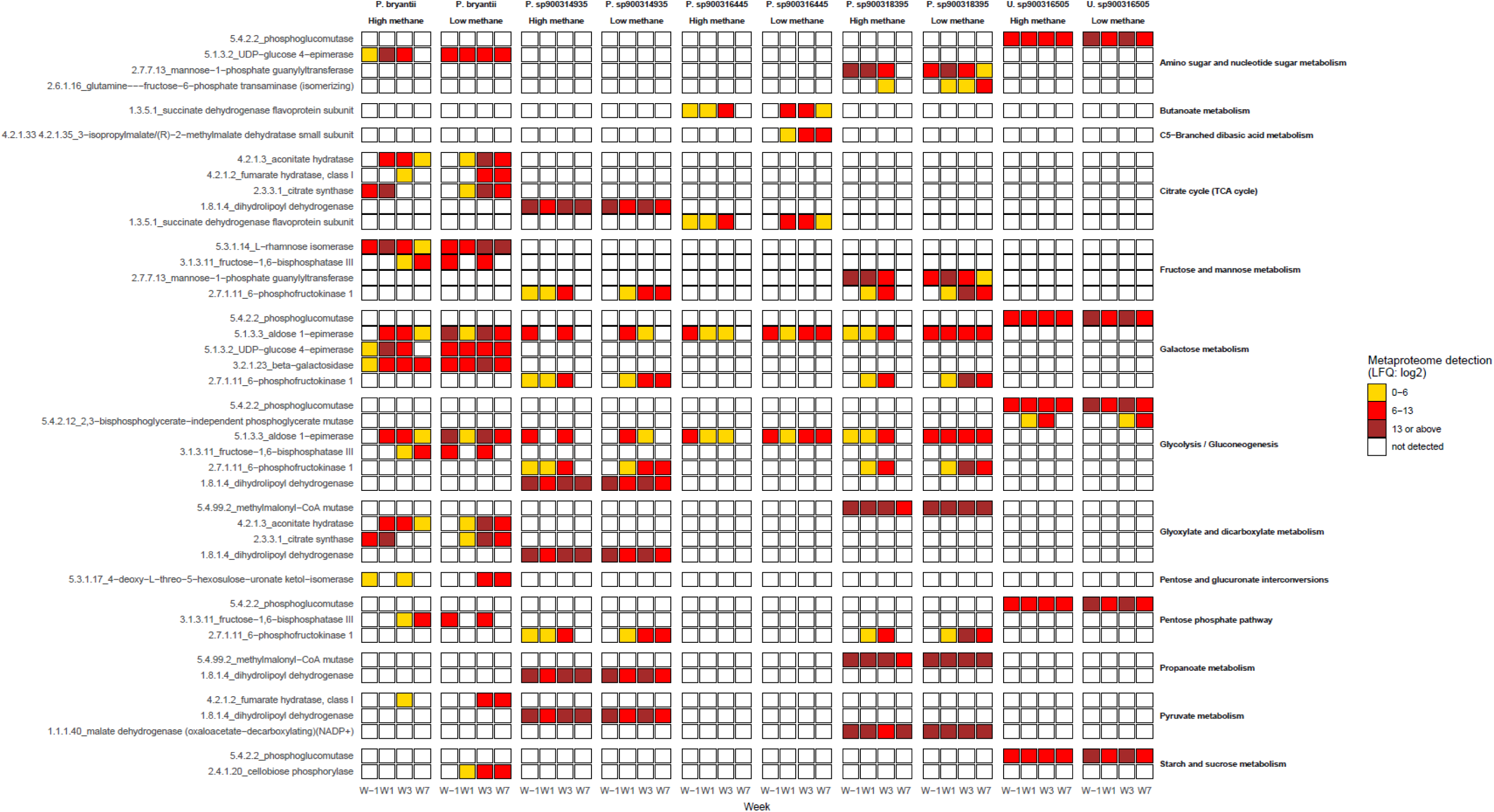
Detected proteins mapped to bacterial metagenome-assembled genomes (MAGs) in the rumen microbiome of low- and high-methane-yielding cows (expressed as g CH_4_/kg dry matter intake). The heatmaps show selected MAGs-*Prevotella bryantii*, *Prevotella* sp. 900314935, *Prevotella* sp. 900316445, *Prevotella* sp. 900318395, and UBA4334 sp. 900316505, with metabolically active proteins represented by EC numbers and enzyme names. Each grid illustrates protein detection across methane phenotypes and sampling weeks, with color intensity indicating protein abundance (white: not detected; yellow: low; orange: medium; red: high; brown: very high).

Expressed proteins and MAG-associated pathways were differential in week 7 (*P* < 0.05). Enzymes involved in carbohydrate and central metabolism, including glycolysis/gluconeogenesis, pentose phosphate, starch and sucrose metabolism, and the citrate and propanoate pathways, were detected throughout the experiment in both phenotypes. The low emitters showed consistently higher enzyme detection in the MAGs *Prevotella bryantii*, *Prevotella sp.* 900316445, and *Prevotella sp.* 900318395 in week 7, except for fructose-bisphosphatase (EC 3.1.3.11). Among the enzymes that can be directly linked to propionate formation, methylmalonyl-CoA mutase (EC 5.4.99.2) and succinate dehydrogenase (EC 1.3.5.1) were detected in several Prevotella MAGs, though the differences in detection between phenotypes were modest and variable across MAGs. In contrast, high emitters exhibited more intermittent detections across several of these pathways. The UBA4334 MAG followed a similar trend, with broader enzyme coverage in low emitters.

## Discussion

By integrating metagenomic and metaproteomic analyses with direct CH_4_ measurements, this pilot study provides initial evidence linking rumen microbiome reconstitution after total rumen content exchange with CH_4_ yield phenotypes in dairy cows. While the small number of animals (n = 2 per phenotype) limits the statistical power and generalizability of the findings, the results are hypothesis-generating and lay the groundwork for larger-scale investigations. This approach extends previous work on rumen microbial resilience [4, 5] by incorporating multi-omics and the CH_4_ dimension, offering novel insights into the host–microbe interactions that govern enteric CH_4_ production.

### Microbial reconstitution and host specificity post-transfaunation

After swapping rumen contents between high and low methane-yielding cows, we observed a striking asymmetry in how the microbial communities re-established. Based on metagenome-derived 16S rRNA gene sequence data, the low CH_4_ emitters gradually reverted to their original microbial community profile, essentially regaining their pre-swap microbiome by weeks 3–7. In contrast, the high CH_4_ emitters did not revert to their original community, and instead, maintained a microbiome resembling that of the low-emitting donor cows. In other words, the high emitters adopted the donor’s microbial community composition and clustered closely with the low-emitter microbiome post-swap. This outcome can be viewed in two ways: either it reflects host specificity in low emitters, or donor community dominance in high emitters. Low emitters’ rumen communities displayed strong resilience and host specificity, quickly recovering their original composition after the swap. This aligns with the findings of Weimer et al. [4], who demonstrated that the rumen microbial community is largely host-specific, with most cows returning to their pre-swap community within 2–8 weeks. Such resilience reflects ecological principles of inertia (resistance to change) and resilience (ability to recover) in the rumen ecosystem. High emitters, on the other hand, did not re-establish their original microbiome and instead retained the donor (low emitter) community. This suggests that the introduced low-emitter microbiota was able to persist in the high-emitting hosts.

Previous complete rumen content swap studies have generally reported a return toward the host’s original microbiota over time. For example, Weimer et al. [4] performed two trials with each one pair of cows and reported 3 out of 4 cows’ communities reverted to pre-swap profiles within 14–62 days. Our finding that 2 out of 4 cows (the high emitters) instead maintained the donor’s community suggests a different reconstitution trajectory, possibly reflecting differences in host factors or methodological approach. We performed a comparatively thorough content swap, including repeated rinsing of both the rumen and omasum to remove indigenous microbes. In contrast, Zhou et al. [5] rinsed only the rumen (not the omasum), and Weimer et al. [4] did not rinse at all, leaving a larger residual of the original microbiota of approximately 5%. The more complete removal of native microbes in our study may have facilitated colonization by the donor microbes in high emitters. Zhou et al. [5] observed in young beef cattle that the individual host has a strong effect on how the microbiome re-establishes, and not all recipients reverted to their original profile within 28 days. In our study with adult cows, we allowed a longer recovery (7 weeks), yet the high emitters still did not regain their original microbes, hinting that host factors “allowed” the donor microbiome to persist in those animals. Since we did not use any chemical treatment to eliminate remaining microbes, we cannot rule out the possibility that microbes in the omasum or those attached to the rumen wall persisted through the washing process and subsequently refaunated the rumen in the low emitters. It is plausible that the low-emitter microbiome was inherently more competitive or better adapted, enabling it to dominate in both its native host and the high-emitter recipients. Overall, our results suggest host specificity in rumen microbiota composition for low emitters (who reasserted their native community), whereas high emitters lacked such strong host imprint, allowing an introduced community to persist.

### Methane emissions remain phenotype-specific and insights from metagenomic and metaproteomic profiles

In our experiment, swapping the microbiota did not change the animals’ methane yield phenotypes. After the swap, the cows that were low emitters remained low emitters. In fact, their CH_4_ yield became even lower, and the high emitters remained high emitters. Our low-emitting cows did not exhibit smaller rumen volumes, contrasting with reports in sheep that link low CH_4_ yield to smaller rumens and shorter retention times [14]. All cows were fitted with rumen cannulas, which can cause minor gas leakage or limited oxygen ingress [15]; however, such effects would be similar across phenotypes and cannot account for the selective CH_4_ decline in low emitters. Both groups showed modest increases in total VFA concentrations and lower acetate-to-propionate ratios by week 7. Although propionate was numerically higher in low emitters, the absolute differences were small, and these fermentation shifts occurred in both phenotypes. Therefore, VFA profiles alone do not provide strong evidence for divergent hydrogen utilization between phenotypes. In other words, the “CH_4_ category” of each cow was unchanged by receiving a different microbiome. This outcome is striking: even though the high emitters ended up hosting a microbial community very similar to that of low emitters, their CH_4_ production stayed high. Similarly, low emitters, after briefly hosting high-emitter microbiota and then reverting to their own, produced as little or less CH_4_ than before. These findings suggest that host factors, rather than the microbial community composition alone, were the dominant drivers of CH_4_ production in these animals.

Methanogenic archaeal abundance and composition, estimated from metagenome-derived 16S rRNA sequences, did not differ significantly between CH_4_ phenotypes at any time point, suggesting that host-side factors rather than methanogen community structure may underline the observed CH_4_ yield differences. Earlier work has shown that methanogens often appear more stable than bacteria following rumen content transfer [6], but our data do not allow conclusions about functional stability or absolute methanogen biomass. More broadly, our results are consistent with earlier work showing that factors beyond the transferred rumen microbiota contribute to CH_4_ yield. For example, Difford et al. [2] reported that host genetic background accounted for approximately 21% of the variation in CH_4_ emissions, whereas the combined effect of rumen bacteria and archaea explained only about 13%. In our study, absolute CH_4_ production, CH_4_ yields and intensity remained close to each cow’s original category despite substantial microbiome turnover after the swap, indicating that replacing the rumen contents alone did not modify the CH_4_ phenotype.

Multi-omics data revealed individualized recovery patterns. Metagenomic and metaproteomic PCAs showed that microbial and functional community structures remained cow-specific and dynamic across time, without a consistent separation by CH_4_ phenotype. Metaproteomic mapping revealed that the enzymes detected in the low emitters were also detected in the high emitters, indicating that the same metabolic functions were present in both phenotypes. When focusing specifically on week 7, several MAGs, including *Prevotella bryantii*, *Prevotella* sp900316445, and *Prevotella* sp900318395, showed higher and more intense protein detection in the low emitters than in the high emitters. The *Prevotella* spp., including *Prevotella bryantii* and *Prevotella* sp900318395, expressed enzymes of the succinate–propionate pathway, including succinate dehydrogenase (EC 1.3.5.1) and methylmalonyl-CoA mutase (EC 5.4.99.2), which are potentially involved in diverting reducing equivalents toward propionate rather than CH_4_ formation [16]. While these enzymes were detected in the low emitters at week 7, the differences between phenotypes were modest and inconsistent across MAGs and thus should be interpreted as suggestive rather than conclusive evidence for metabolic hydrogen rerouting. Across the broader metaproteome, nearly all MAGs, except *Prevotella* sp900316445, also expressed enzymes involved in glycolysis, gluconeogenesis, and the pentose phosphate pathway, suggesting that carbohydrate metabolism remained active while energy flow was redirected toward succinate and propionate production. Such rerouting of carbon and electron flux is generally associated with reduced CH_4_ output, since hydrogen generated during carbohydrate fermentation is internally metabolized instead of being transferred to methanogens [7]. However, the overall intensity patterns in week 7 did not show a clear or consistent separation between low and high emitters, and the observed differences varied by MAG and by enzyme. Tricarboxylic acid (TCA) cycle enzymes such as citrate synthase (EC 2.3.3.1) and fumarate hydratase (EC 4.2.1.2) were detected more frequently and with higher intensity in the low emitters in weeks 3 and 7. These enzymes support NADH reoxidation within the TCA cycle, providing an additional sink for reducing power and helping to limit hydrogen accumulation. The metaproteomic data are consistent with but do not conclusively demonstrate metabolic rerouting in low-methane cows. The detection of succinate–propionate pathway enzymes in Prevotella-dominated communities of low emitters raises the possibility that alternative hydrogen utilization contributes to their lower CH_4_ emissions, but these observations require validation in larger studies with greater statistical power. We note that automated pathway assignment tools can sometimes assign enzymes to pathways where they are not the primary catalysts. In this study, we focused on enzymes with well-established roles in their assigned pathways and acknowledge that some detected enzymes participate in multiple metabolic routes. Metaproteomic coverage was not comprehensive for all central metabolic pathways; for instance, glycolytic enzymes were detected in fewer MAGs than expected given the carbohydrate-rich diet. The absence of certain enzymes should therefore not be taken as evidence that the corresponding pathways are inactive.

## Conclusion

Complete rumen content swap between low- and high-methane-yielding cows did not alter their CH_4_ phenotype, suggesting that host factors may play an important role in determining CH_4_ production. Low emitters re-established their original microbial community and maintained low CH_4_ yield, whereas high emitters sustained high emissions despite hosting donor-like microbiota. Low emitters further reduced their CH_4_ yield despite receiving high-emitter microbiota, and multi-omics profiling suggested higher activity of Prevotella-associated propionate pathway enzymes, although these differences were modest and require confirmation. The findings suggest that durable CH_4_ mitigation will require integrating host selection with strategies that enhance microbial hydrogen utilization.

## Material and methods

The animal experiment was carried out at the Livestock Production Research Center and the Metabolism Unit of the Norwegian University of Life Sciences. The experimental procedures complied with the Norwegian legislation of animal welfare and were approved by the Norwegian Food Safety Authority (Mattilsynet; FOTS ID: 24379).

The experiment began with the screening of 20 Norwegian Red dairy cows over a 30-day period to record feed intake, milk yield, body weight (BW), and CH_4_ emissions using the GreenFeed system (C-Lock Inc., Rapid City, SD, USA). Based on their CH_4_ yield, two low- and two high-emitting cows were selected for rumen cannulation, which was performed approximately eight weeks before their expected calving date. Two months after calving, a complete rumen content swap was conducted, including total rumen evacuation, thorough washing of the rumen and omasum, and reintroduction of donor rumen content. Rumen content samples were collected at eight time points during the subsequent two months for metagenomic and metaproteomic analyses of microbial (bacteria, protozoa, archaea) composition and function. Methane emissions were measured during the screening phase in the first lactation (pre-cannulation) and again at the end of the experimental period, approximately two months after the rumen content swap, to evaluate potential changes in CH_4_ yield relative to the pre-swap baseline (Fig. 6).

**Fig. 6.**
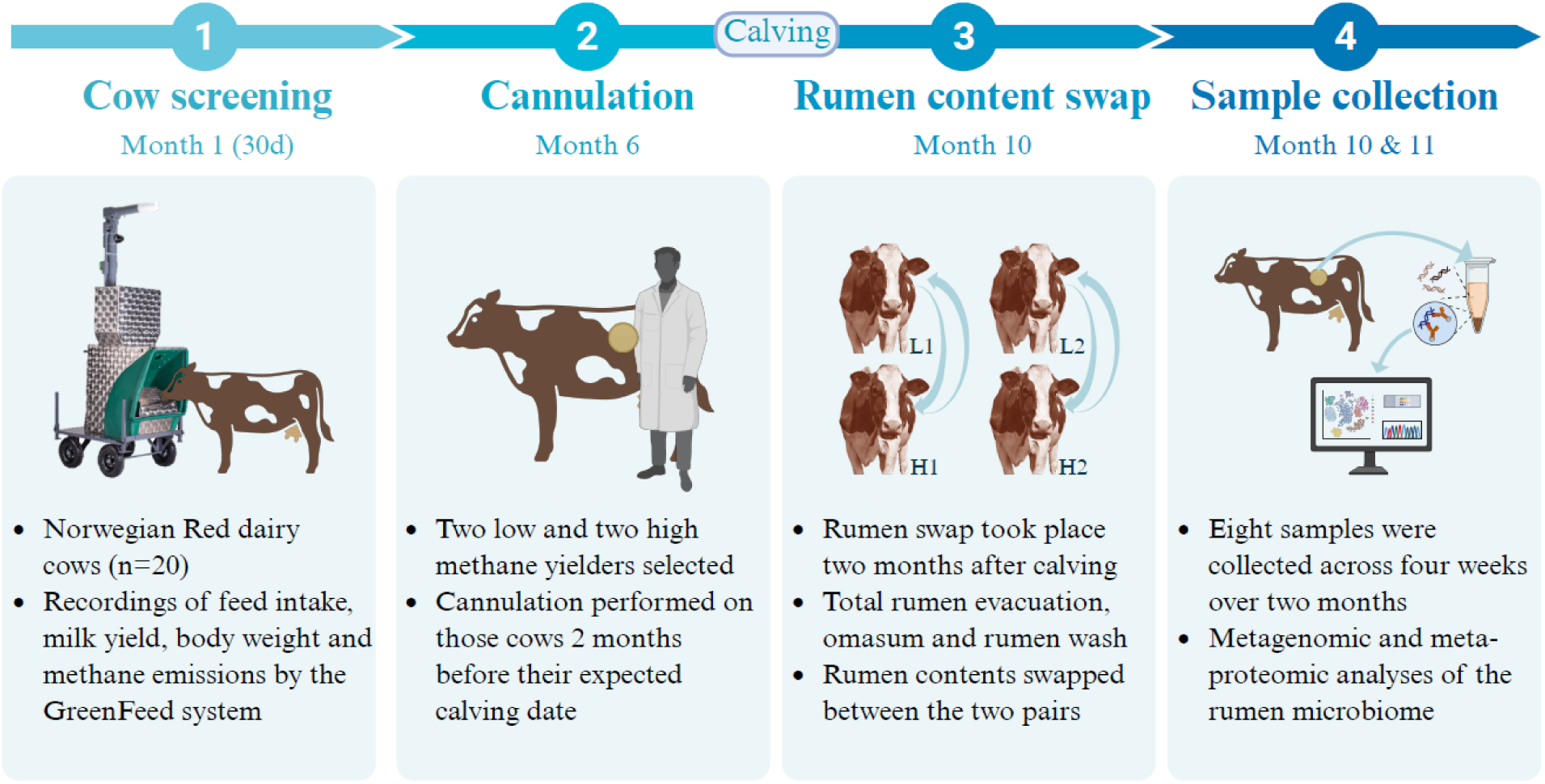
Study workflow: (1) 20 cows were screened for feed intake, milk yield, body weight, and methane emissions for a month; (2) The selected primiparous cows were cannulated 2 months pre- calving; (3) The rumen contents of low methane yielders were swapped with those of the high methane yielders 2 months post-calving (methane yield is expressed as g CH_4_/kg dry matter intake); (4) Rumen samples were collected at eight time points across four weeks (week −1, one week before the swap; and weeks 1, 3, and 7, one, three, and seven weeks after the rumen content swap) over a two-month period for metagenomic and metaproteomic analyses of microbial composition and function.

### Animals and housing

The screening of 20 Norwegian Red dairy cows for CH_4_ yield was conducted at the Livestock Production Research Center (Norwegian University of Life Sciences, Ås, Norway) as part of a larger experiment involving 60 cows described by Álvarez et al. [10]. The cows were housed in a free-stall barn with concrete floors and rubber mats topped with sawdust. Of the 60 cows, 34 with valid CH_4_ data were available for CH_4_ emission ranking. Based on their calving dates and the culling plan that aligned with the rumen swap experiment schedule, 20 cows were screened to identify four cows for the rumen cannulation and subsequent rumen content swap.

At the start of the screening (Fig. 1), the 20 cows averaged 79 ± 30 days in milk, 35 ± 8 kg/day milk yield, and 611 ± 66 kg BW; six were primiparous and fourteen multiparous (second to fourth lactation). The rumen content swap was later conducted in the tie-stall barn of the Livestock Production Research Center. The four cows selected for this experiment were all primiparous and, at the start of the rumen swap, averaged 49 ± 20 days in milk, 32 ± 4 kg/day milk yield, 602 ± 36 kg BW, and 24 ± 2 kg/day DMI. The tie-stall was equipped with rubber mats and sawdust bedding, and all cows had *ad libitum* access to water.

### Diet, feeding, and milking

During the screening period, cows were fed a common grass silage *ad libitum* (5–10% refusals of the offered amount) and a commercial concentrate provided according to milk yield, as described by Álvarez et al. [10]. The silage was prepared in round bales containing, on a seed-weight basis, 50% timothy (*Phleum pratense*), 25% meadow fescue (*Festuca pratensis*), 15% meadow grass (*Poa pratensis*), and 10% white clover (*Trifolium repens*). Equal proportions of early- and late-harvested silage were mixed using a Siloking Duo 1814 mixer (Kverneland, Norway) and offered once daily at 07:00 h through 40 individual automatic feeders (BioControl AS, Pakkestad, Norway) equipped with vertically moving gates. Feed intake per cow was recorded automatically via electronic identification, and refusals were removed before the next feeding.

The concentrate consisted (in descending proportion) of barley, soybean meal, oats, maize, wheat, sugar beet pellets, molasses, rapeseed cake, beans, and a mineral–vitamin premix. One kilogram per day was provided in the GreenFeed units, while the remainder was distributed via the automatic milking robot (DeLaval International AB, Tumba, Sweden) and three concentrate feeders. Each feeding event supplied up to 4 kg concentrate with a minimum 4-h interval between feedings. Chemical compositions of the silage and concentrate are reported by Álvarez et al. [10]. Briefly, on a dry matter (DM) basis, early- and late-cut silages contained 254, 468, and 215 g/kg and 237, 597, and 144 g/kg of DM, NDF, and CP, respectively, while the concentrate contained 861, 180, and 198 g/kg. Feed samples were oven-dried at 103 °C for 24 h to constant weight to determine DM content.

During the rumen content swap experiment, cows were offered a timothy-based grass silage mixture ad libitum (10% refusals of the offered amount) and a commercial concentrate (Drøv Fase 1, Norgesfor, Norway) according to milk yield. The silage was produced from a third-year ley (SPIRE Surfôr/Beite Pluss 10; Felleskjøpet Rogaland Agder, Stavanger, Norway) containing 42% timothy, 23% meadow fescue, 15% blue grass, 10% white clover, and 10% perennial ryegrass. Silage was provided in feeding troughs, while the concentrate was offered in buckets placed within the same trough. The daily allowance was divided into three equal feedings at 07:00, 13:00, and 19:00 h, and refusals were collected before morning feeding. Except for DM, which was used to calculate DMI, the chemical composition of feeds during this phase was not analyzed. All cows had ad libitum access to water throughout both experimental phases.

Milk yield was automatically recorded at each milking by the automatic milking system (AMS; DeLaval International AB) during the screening period and by a manual milking machine (VEVOR 25 L, Rancho Cucamonga, CA, USA) at 07:00 and 19:00 h during the rumen swap experiment. Milk samples were collected only during the screening period, over a 48-h period at the end of week 4, preserved with Bronopol, and stored at 4 °C until analysis for fat, protein, and lactose using Fourier Transform Infrared (FTIR) spectroscopy (Bentley FTS/FCM or Combi 150; Bentley Instruments Inc., Chaska, MN, USA) at the TINE laboratory (Trondheim, Norway). Body weight was automatically recorded after each milking during screening using an electronic scale at the AMS exit and over a 7-day period before screening began. During the rumen swap experiment, cows were weighed once at the beginning and once at the end using a portable electronic scale, and the average value was used for analysis.

### Methane measurement

Methane production was measured during the 30-day screening period and again for seven consecutive days at the end of the rumen content swap experiment, approximately two months after the swap. Methane data were also collected immediately before the swap, but these measurements could not be included due to technical issues affecting the results of one animal.

During the screening period, CH_4_ production was recorded using two GreenFeed systems (C-Lock Inc., Rapid City, SD, USA) installed in the free-stall barn, which the cows visited voluntarily. The GreenFeed system performed spot measurements of gases exhaled by individual cows, identified by a radio frequency identification (RFID) ear tag, during visits to the feeding hood. Each unit was fitted with one hood dispensing pelleted concentrate to attract the animals. The GreenFeed units dispensed five drops of 40 g concentrate every 25 seconds per visit, with a minimum interval of 4 hours between successive visits. A measurement was considered valid when the animal’s head remained correctly positioned within the hood for at least two minutes. On average, cows visited the GreenFeed systems 3.2 times per day.

Methane concentrations were determined using nondispersive infrared (NDIR) sensors that measured background gas concentration, differential gas accumulation during the animal’s visit, and sensor calibration coefficients. Automated calibration was performed using nitrogen, carbon dioxide, and CH_4_ reference gases to verify sensor response. A gravimetrically measured amount of carbon dioxide gas was released at the feed position to determine gas recovery manually at the beginning and end of data collection. The GreenFeed data were downloaded and managed through the proprietary web-based system provided by C-Lock Inc. During the screening, a viral infection in week 3 caused a temporary reduction in milk yield across all cows; therefore, data from that period were excluded. Consequently, the number of valid observation days per animal was reduced from 30 to 20.

### Ruminal and omasal evacuation, washing and rumen content swap

Rumen evacuation was performed manually by first removing wet solids through the cannula and subsequently collecting residual ruminal fluid using a 0.25-L plastic cup. During the procedure, the experimenters wore shoulder-length polyethylene gloves, and face masks and protective face shields due to pandemic control measures. The removed contents were placed in pre-cleaned plastic containers pre-warmed to 39 °C in a water bath to maintain microbial viability. Ruminal contents were separated into solid and liquid fractions by hand-pressing the material through a sieve bucket. Both fractions were weighed, and according to the liquid-to-solid ratio, a 500 mL representative sample of homogenized rumen content was collected and oven-dried at 103 °C for DM determination. The DM value of this composite was used to calculate the total DM of the rumen contents.

The omasum was emptied by repeatedly washing with warm tap water (approximately 39 °C) using a rubber hose (10 mm outer diameter) equipped with an electronically powered spiral drill. One end of the hose was immersed in a water-filled bucket to provide a continuous flow of warm water. The mixture of wash water and omasal particulate matter that returned to the rumen through the reticulo-omasal orifice was removed by suction and discarded. The process was repeated several times, using approximately 10 L of water for each rinse, followed by two additional washings of the reticulo-rumen. A video of the washed rumen can be found in the supplemental material (https://doi.org/10.18710/EU13P5).

After evacuation and washing of both the rumen and omasum, the rumen contents (solids and liquids) from donor and recipient cows were swapped. The wet solids were manually inserted into the reticulo-rumen of the recipient cow through the cannula, followed by the addition of the corresponding liquid fraction. The entire evacuation, washing, and content swap procedure between the two low- and two high-methane-emitting cows was completed within approximately one hour.

### Rumen content sampling

Rumen content (liquid and solid) was collected over a two-month period at eight time points corresponding to four experimental weeks: week −1 (days −5 and 0, pre-swap), week 1 (days 2 and 7), week 3 (days 20 and 24), and week 7 (days 48 and 52) (Table S1). Duplicate sampling at each of these weeks captured the major stages of the microbial intervention, including pre-swap, early, mid, and late post-swap.

Samples were collected two hours after feeding (09:00 h) from the central region of the rumen using a plastic cup. A portion of each sample was transferred into 10 mL tubes pre-filled with 0.5 mL of concentrated (98–100%) formic acid for analysis of volatile fatty acids (VFA) and ammonia and stored at 4 °C until analysis. Another portion was placed into 2 mL Eppendorf tubes containing 0.9 mL RNAlater for metagenomic analysis, kept at 4 °C overnight, and then stored at −80 °C. Additionally, 2 mL of rumen content was transferred into Eppendorf tubes without RNAlater, snap-frozen in liquid nitrogen, and stored at −80 °C for metaproteomic analysis. A subsample was used immediately for pH measurement, as described below, and the remaining material was returned to the rumen. All samples were collected in duplicate for each cow at each time point.

### Analysis of fermentation parameters

Ruminal pH was measured at each sampling using a pH meter (WTW 3320; Weilheim, Germany) equipped with a pH sensor (Polyplast Pro; Hamilton Bonaduz AG, Switzerland) that was calibrated at pH 7.00 and 4.00 immediately before use. Volatile fatty acids were analyzed by gas chromatography (TRACE 1300; Thermo Fisher Scientific S.p.A., Milan, Italy) equipped with a Stabiliwax-DA column (30 m × 0.25 mm i.d., 0.25 μm film thickness). Ammonia was determined according to AOAC Official Method 2001.11 (AOAC International, 1995), as described by Thiex et al. [11], using a Kjeltec 8400 analyzer (Foss Analytical, Hillerød, Denmark).

### Genome-centric metagenomic analysis

The metagenomic and bioinformatic analyses were performed by DNAsense ApS (now Cmbio, Aalborg, Denmark).

### DNA extraction

Sample biomass was homogenized and aliquoted for optional multiple extraction rounds and keeping a reserve. DNA extraction was initially done using the standard protocol for RNeasy PowerMicrobiome Kit (Qiagen, Germany) with minor modifications: custom reagent volumes were used, PM4 buffer was replaced with 70 % ethanol in initial extraction mix, and bead beating was performed at 6 m/s for 4×40s. Before DNase treatment with the DNase Max kit (Qiagen, Germany), the extracts were split in two with one half left untreated such that both an RNA and DNA preparation were obtained. Gel electrophoresis on the Tapestation 2200 (Agilent, USA) was used to validate product size and purity of nucleotide extracts, and concentration measurements were done using Qubit (Thermo Fisher Scientific, USA). RNA screentapes and the Qubit RNA HS assay kit was used for RNA extracts, and for DNA extracts the D1000 screentapes and Qubit dsDNA HS assay kit. The extracted DNA was used for preparing Illumina metagenome sequencing libraries, and the RNA was stored for potential later use. Oxford Nanopore Technologies (ONT) sequencing has high requirements for DNA input and integrity, and the DNA from the initial extraction round was deemed of suboptimal quality for ONT. A second round of DNA extraction was done using the DNeasy Power Soil kit following the manufacturer’s recommendations (Qiagen, Germany) with minor modifications. The Circulomics Short Read Eliminator XS Kit was used for DNA fragment size selection (Circulomics, USA). Gel electrophoresis using Tapestation 2200 (Genomic DNA and D1000 screentapes, Agilent, USA) was used to validate product size and purity of a subset of DNA extracts. DNA concentration was measured using both Nanodrop and Qubit dsDNA HS Assay kit (Thermo Fisher Scientific, USA). The extracted DNA was used for preparing ONT sequencing libraries.

### Illumina sequencing and Nanopore sequencing

The DNA was quantified using Qubit (Thermo Fisher Scientific, USA) and fragmented to approximately 550 bp using a Covaris M220 with microTUBE AFA Fiber screw tubes and the settings: duty factor 10%, peak/displayed power 75W, cycles/burst 200, duration 40s and temperature 20°C. The fragmented DNA was used for metagenome preparation using the NEB Next Ultra II DNA library preparation kit. The DNA library was paired-end sequenced (2 x 150 bp) on a NovaSeq S4 system (Illumina, USA). The reads generated were approximately 150 million for 32 rumen samples. Four long-read sequencing libraries (one per animal) were prepared according to the SQK-LSK110 protocol (ONT, Oxford, United Kingdom), generating 15−20 Gb data per sample. The N50 was 5 to 10 kb. Approximately 100 fmole was loaded onto primed FLO-MIN106D (R9.4.1) flow cells and sequenced on the GridION platform using MinKNOW Release 21.05.12.

### Data preprocessing

Raw Illumina reads were filtered for PhiX using Usearch11 and trimmed for adapters using cutadapt (v. 2.8). Raw Oxford Nanopore Fast5 files were basecalled in Guppy v. 5.0.12 using the highaccuracy (hac) algorithm. Basecalled fastq data was then adapter-trimmed in Porechop v. 0.2.4 Porechop using default settings. NanoPlot v.1.27 was used to assess quality parameters of the basecalled data. The trimmed data was then filtered in Filtlong v. 0.2.0 with min_length set to 1000 bp and min_mean_q set to 90.

### Taxonomic classification of metagenomic reads

Forward trimmed Illumina reads were mapped to the SILVA 138 SSU database [12] using minimap2 (v. 2.22-r1101). Mappings were further processed via RStudio IDE (4.1.0 (2021-05-18)) running R version 4.1.1 (2021-08-10) and using the R packages: ampvis (2.7.17), tidyverse (1.3.1). The term operational taxonomic unit (OTU) is used to describe the metagenome derived 16S rRNA gene sequences. However, sequences were treated as zero-radius OTUs (no clustering), which can also be recognized as assigned sequence variants (ASV).

### Metagenome assembly and polishing

Draft metagenomes were assembled separately for each of the four ONT libraries with Flye (v.2.9) by setting the metagenome parameter (–meta). The assembled metagenomes were subsequently polished with quality-filtered ONT data using one round of polishing with Minimap2 (v. 2.22-r1101) and Racon (v.1.4.13) and two rounds of polishing with Medaka (v.1.4.3). The metagenome was finally polished with minimap2 (v. 2.22-r1101) and racon (v.1.4.13) using respective Illumina data. The metagenomes were visualized in RStudio IDE (4.1.0 (2021-05-18)) running R version 4.1.1 (2021-08-10) and using the mmgenome2 R package (v. 2.1.3).

### Binning and classification

Each of the four metagenome assemblies was subjected to independent and automated genome binning (minimum length set to 500 Kbp) using Metabat2 (v. 2.12.1), Maxbin2 (v. 2.2.7) and Vamb (v. 3.0.2). The metagenome assembled genomes (MAGs) from each holometagenome were subsequently dereplicated using Das Tool (v. 1.1.3). The best-scoring MAGs from each holometagenome were pooled and dereplicated using dRep (v. 3.2.2) and default settings. MAG completeness and contamination were assessed using CheckM (v. 1.1.2). Dereplicated MAG abundance was calculated using CoverM (v. 0.6.1).

MAGs were taxonomically classified using the Genome Taxonomy Database toolkit (v. 1.5.1). 16S rRNA and tRNA gene sequences were extracted and identified using Barrnap (v. 0.9) and tRNAscan-SE (v. 2.0.5), respectively. Dereplicated MAGs were annotated using Prokka (v. 1.14.6). The final, dereplicated data contained 309 MAGs, out of which 230 had 16S rRNA genes present.

### Metaproteomic data generation and analysis

Metaproteomic analyses were conducted on all 32 rumen content samples collected from the four cows. For each sample, 300 µL of rumen fluid was mixed with 150 µL lysis buffer (30 mM dithiothreitol, 150 mM Tris-HCl, pH 8.0, 0.3% Triton X-100, and 12% SDS) and 4 mm glass beads (≤160 µm). Samples were briefly vortexed, incubated on ice for 30 min, and lysed using a FastPrep-24 Classic Grinder (MP Biomedical, OH, USA) for 3 × 60 s at 4.0 m/s. The lysates were centrifuged at 16,000 × g for 15 min at 4 °C, and the supernatant was collected. Protein concentration was determined using the Bio-Rad DC Protein Assay (Bio-Rad, Hercules, CA, USA) with bovine serum albumin as the standard. Absorbance was measured at 750 nm using a BioTek Synergy H4 Hybrid Microplate Reader (Thermo Fisher Scientific, Waltham, MA, USA).

Aliquots containing 40–50 µg of total protein were prepared in SDS buffer, heated at 99 °C for 5 min, and separated by SDS–PAGE using Any-kD Mini-PROTEAN TGX Stain-Free gels (Bio-Rad, Hercules, CA, USA) in a short 2-minute run for sample clean-up. The gel was stained with Coomassie Brilliant Blue R-250, and visible bands were excised and cut into 1 × 1 mm pieces for reduction, alkylation, and trypsin digestion. Peptides were purified using C18 ZipTips (Merck Millipore, Darmstadt, Germany), dried, and analyzed by nano-LC-MS/MS on a Q-Exactive Hybrid Quadrupole-Orbitrap mass spectrometer (Thermo Fisher Scientific, Waltham, MA, USA).

Mass spectrometry raw data were processed with FragPipe (v.19.0) and searched against a sample-specific protein sequence database containing the 309 metagenome-assembled genomes generated in this study, using MSFragger. The database was supplemented with common contaminant proteins (e.g., human keratin, trypsin, bovine serum albumin) and reversed sequences of all protein entries for estimation of false discovery rates (FDR). Variable modifications included oxidation of methionine and N-terminal acetylation; carbamidomethylation of cysteine was used as a fixed modification. Trypsin was set as the digestive enzyme, with a maximum of one missed cleavage and mass tolerances of 20 ppm for both precursor and fragment ions.

Results were filtered to 1% FDR, and quantification was performed with IonQuant using normalization across samples and the “match between runs” feature to increase peptide identification. Protein groups were retained if quantified in at least 50% of replicates within one CH_4_ phenotype. Potential contaminant proteins were excluded. Summed protein abundances (label-free quantification, LFQ values) were calculated for each MAG following the workflow of Kleiner et al. [13], and differences between low- and high-methane phenotypes were assessed using unpaired two-sided Student’s t-tests (*P* < 0.05). After filtering, metaproteomic analysis identified 21410 unique protein groups across the 32 samples, with functional assignment linked to the 309 MAGs reconstructed from the same animals’ rumen metagenomes.

### Statistical analysis

All statistical analyses were conducted in R (version 4.3.0; R Core Team, 2023). Differences were considered statistically significant at *P* < 0.05, and results are reported as mean ± standard deviation unless stated otherwise. Descriptive statistics were used to summarize feed intake, milk yield, BW, and CH_4_ emission data from the screening period for the 20 cows and for the four cows selected for the rumen content swap.

Rumen fermentation parameters (pH, VFA profiles, and ammonia) were summarized descriptively to illustrate temporal changes across sampling weeks within each methane phenotype. No statistical comparisons were conducted for these parameters.

Microbial community composition was assessed using principal component analysis (PCA) implemented in the ampvis2 package (v. 2.7.27) to compare overall bacterial and archaeal profiles from metagenome-derived 16S rRNA gene sequence data. PCA was also performed on MAG abundance data and metaproteomic profiles to visualize microbial and functional shifts following the rumen content swap. Differentially abundant MAGs and taxa were identified in samples from week 7 using unpaired two-sided Student’s t-tests (*P* < 0.05).

For metaproteomic data, protein quantification was based on summed label-free quantification (LFQ) intensities across biological replicates within each CH_4_ phenotype. Proteins assigned to MAGs were further annotated to KEGG pathways, and pathway-specific enzyme abundance profiles were visualized as heatmaps. Differentially expressed proteins and MAG-associated pathways were identified using unpaired t-tests (*P* < 0.05).

## Declarations

### Ethics approval and consent to participate

All procedures involving animals complied with the Norwegian legislation of animal welfare and were approved by the Norwegian Food Safety Authority (Mattilsynet; FOTS ID: 24379). The study was conducted and reported in accordance with the ARRIVE guidelines for reporting animal research.

### Consent for publication

Not applicable.

### Availability of data and material

The shotgun metagenomic sequencing data (Illumina short-read) generated in this study have been deposited in the NCBI Sequence Read Archive (SRA) under the BioProject accession PRJNA1405063. Raw mass spectrometry–based proteomics data have been deposited in the MassIVE repository under accession MSV000100725 (DOI: 10.25345/C5ZW1960Z).

### Author contributions

P. Niu: Formal analysis, investigation, data curation, visualization, writing – original draft, and writing – review & editing; C. Kobel: Investigation, data curation, and writing – review & editing; V. Aho: Data curation and writing – review & editing; C. Álvarez: Resources and writing – review & editing; E. Prestløkken: Methodology and writing – review & editing; P. Lund: Methodology and writing – review & editing; P.B. Pope: Methodology, supervision, data curation, and writing – review & editing; A. Schwarm: Conceptualization, funding acquisition, methodology, validation, supervision, data curation, and writing – review & editing.

## Acknowledgement

The authors thank the staff at the Livestock Production Research Center and the Metabolism Unit for their assistance with rumen cannulation, silage production, and animal care and sampling. We also thank the laboratory staff at the Department of Animal and Aquacultural Sciences for their support with sample preparation and chemical analyses.

